# Ant community potential for pest control in olive groves: management and landscape effects

**DOI:** 10.1101/2020.05.18.103028

**Authors:** Carlos Martínez-Núñez, Pedro J. Rey, Teresa Salido, Antonio J. Manzaneda, Francisco M. Camacho, Jorge Isla

## Abstract

Ants are important pest control agents in many agroecosystems worldwide. However, little is known about how management, ecological contrast generated by different agricultural practices, and landscape complexity affect their communities and the potential pest control that they can provide.

Here, we surveyed ant communities in 40 paired olive groves with different ground-herb cover management across 20 localities spanning a wide range of landscape complexity at the regional scale. We also conducted experiments with dummy plasticine models in 18 of these groves to explore the ant potential to control the two main pests of the olive tree (olive moth and olive fly). In addition, we calculated an index, the Ant-community Predation Function (APF), which informs about the predation pressure exerted by ant communities over these pests.

Results show that: a) extensive management at intermediate landscape complexity maximizes the abundance and pest control by ants; b) the ecological contrast affects ant abundance and ant richness but does not impact the predation function; c) APF is a good measure of pest control potential at the community level; and, d) *Tapinoma nigerrimum* is an important ant species for pest control in our system, which seems little affected by local or landscape heterogeneity.

This study advances our knowledge about local management and landscape effects on ants and their potential contribution to pest control in olive groves. Extensive herb cover management and landscape heterogeneity increased ant predation pressure in olive groves.

## Introduction

Ants are one of the most abundant, diverse, and widespread insect families in permanent agroecosystems (e.g. Philpott & Armbrecht, 2006; Rey et al., 2019). Many ant species are active predators with high energetic demands, which confers them a strong potential as pest control agents (Thurman, Northfield, & Snyder, 2019; Way & Khoo, 1992). Their effectiveness for biocontrol, and for yield increase, has proven important in many systems (mainly tropical), including groves, agroforests and arable crops (reviewed by Offenberg, 2015). Some notable cases include the overall positive effect of weaver ants in tropical tree crops (Thurman et al., 2019), or the biocontrol provided by *Azteca sericeasur* and other ant species against the coffee borer (Escobar-Ramírez, Grass, Armbrecht, & Tscharntke, 2019; Morris, Vandermeer, & Perfecto, 2015).

Habitat management in the form of agricultural extensification, or ecological intensification (*sensu* Bommarco, Kleijn, & Potts, 2013), arises as a sustainable alternative to increase the numbers of natural enemies, such as ants, and reduce the use of pesticides against pest species (Gurr, Wratten, Landis, & You, 2017). However, current trends of intensive agriculture and landscape simplification can affect negatively these and other arthropod natural enemies, threatening their communities, and reducing their pest control potential (Aristizábal & Metzger, 2019; Gallé, Happe, Baillod, Tscharntke, & Batáry, 2019; Haan, Zhang, & Landis, 2019; Milligan, Johnson, Garfinkel, Smith, & Njoroge, 2016). Nonetheless, some ant species can tolerate environmental disturbances, and most studies measuring ant-mediated pest control have been conducted in tropical agroecosystems. Therefore, it remains uncertain to what extent management and landscape complexity can affect ant communities and their potential pest control in dryland agroecosystems, such as Mediterranean olive groves, the more extended woody crop across the Mediterranean basin.

Several authors have characterized the soil arthropod fauna in olive groves (Gonçalves & Pereira, 2012) and have explored management (Cotes, Campos, Pascual, García, & Ruano, 2010; Ruano et al., 2004) and environmental heterogeneity (Gkisakis, Volakakis, Kollaros, Bàrberi, & Kabourakis, 2016) effects on epigeal insect species and pest control. For instance, Picchi et al., (2017) found higher predation rates on the soil for the olive moth (an overwintering pest) in organically managed olive groves. However, although ants are one of the most abundant soil arthropods in olive groves (Santos, Cabanas, & Pereira, 2007), and some species, such as *Tapinoma nigerrimum* have been pointed as an important species for biocontrol (Morris et al., 2002), the ant pest control function remains essentially understudied in this agroecosystem. Moreover, very few studies have focused on the main drivers of ant communities. Two studies explore local management and landscape effects on ant abundance (Gkisakis et al., 2016) and diversity (Rey et al., 2019) in olive groves. The former, found no differences in abundance due to management or landscape complexity. The latter, conducted at the regional scale, showed how both ground herb cover management intensification and landscape simplification reduced ant species richness on olive farms. Although ants predate on the olive moth (Morris et al., 2002; Morris et al., 1998) and the olive fly pupae (Orsini, Daane, Sime, & Nelson, 2007), there are still scarce evidences about their real potential for pest control at the community level. Even less is known about how management intensification, landscape homogeneity or ecological contrast (i.e. the degree of environmental contrast produced by differing agricultural management regimes) can affect ant communities and their pest control function in olive groves.

In this study, we focused on these issues using a large-scale survey and an experimental approach. We conducted ant surveys on 40 olive farms to characterize the ant communities at the farm scale. These surveys were followed by field experiments to calculate the Ant-community Predation Function index (hereafter, APF), using fresh prey offerings to ants, to characterize ant-specific response to pests. This index accounts for the quantitative (abundance) and qualitative (appetence) components of predation, and represents the potential predation pressure that the local ant community can exert on the olive moth and the olive fly (Martínez-Núñez & Rey, 2020a) and it thus is a proxy of pest control potential. We also carried out artificial dummy model prey experiments on 18 farms to assess ant predation pressure, and compare ant attack rates across farms with different herb cover management (intensive vs. extensive). Finally, we analyzed how the interplay between agricultural management (considering different levels of ecological contrast) and landscape simplification can drive differences in ant communities, and APF. Based on sound hypotheses in agroecology, such as the intermediate landscape complexity hypothesis (Tscharntke et al., 2012) and the ecological contrast hypothesis (Kleijn et al., 2011; Marja et al., 2019), we predict that: 1) ant abundance and APF will be higher on extensively managed farms and complex landscapes; 2) the magnitude of the differences between farms with different management (i.e,. the effectiveness of the agriculture extensification to recover ant-driven pest control) will be landscape context dependent (Tscharntke et al., 2012); 3) pairs of farms with higher ecological contrast will show stronger differences in comparison to pairs with lower ecological contrast (Kleijn et al., 2011); 4) ant’s attack rates to dummy models will be higher in extensively managed farms; and 5) benefits of agri-environmental management regimes on ant abundance will parallel benefits on APF and attack rates. Finally, we also expect that; 6) the generalist species *Tapinoma nigerrimum* will account for an important contribution to APF and will not be affected by local management nor landscape simplification.

## Materials and Methods

### Study design and study system

This study was conducted in 40 paired olive farms, sited in 20 localities across Andalusia, Spain (see Rey et al. 2019 for details on localities), the region that harbors the highest number of olive trees worldwide. Studied olive farms covered altogether an area of ca. 3.000 ha, and represented an extent of ca. 18.142 Km^2^, encompassing a wide range of landscapes, from very simple landscapes highly dominated by olive groves, to complex montane ecosystems with some olive farms embedded on them (Rey et al., 2019). In each locality, a pair of relatively close olive farms was selected. The pair of farms in each location shared the same landscape context, but differed notably in management. One of the farms had an intensive herb cover management, preventing the growth of herb cover for the most part of the year. In contrast, the herb cover on the other farm was managed extensively (i.e. maintaining the herb cover throughout the year). In addition to this intensive-extensive dichotomy, in ten out of the 20 localities, the extensively-managed olive grove was also under organic management (i.e. without application of synthetic pesticides or fertilizers), adding a further step for ecological contrast (sensu Kleijn et al., 2011; Scheper et al., 2013) between the farms of these ten pairs. This higher ecological contrast also translated into stronger quantitative differences in herb richness and herb cover (Martínez-Núñez et al. 2020b). This design allowed to test the interacting effects of landscape complexity and management on the predation pressure exerted by ants in olive groves. In addition, the two subsets of farms with differing ecological contrast (ten pairs with low contrast and ten pairs with high contrast), permitted us to test the effects of ecological contrast on the ant community and their potential for pest control.

To study the role of ants as pest control agents, we focused on their effect against the two main pest species of the olive tree, the olive moth (*Prays oleae*, Bernard 1788; Yponomeutidae) and the olive fly (*Bactrocera oleae*, Rossi 1790; Tephritidae), which can cause important yield loss (Malheiro, Casal, Baptista, & Pereira, 2015; Paredes et al., 2019). Both pest species have a complex life cycle, closely matching the phenology of the olive tree. The olive moth has three generations, the phillophagous larva, that feeds on the leaves in winter, the anthophagous larva, which develops in spring, feeding on the olive floral buttons and flowers (this larva is highly exposed and probably the most vulnerable to the predation by ants foraging on the trees), and the carpophagous larva (feeding inside the fruit after falling to the ground during autumn) (Pelekassis, 1962). The olive fly is vulnerable to ants at the end of autumn, when larvae finish feeding and growing inside the olive fruits and fall down to the ground to pupae and overwinter (Daane & Johnson, 2010).

### Landscape complexity

We measured the compositional and configurational heterogeneity of the landscapes in a buffer of 2 km radius around the centroid of each pair of farms (N=20). We approached landscape complexity as a multivariate and non-simplistic measure, which represents better its functional and intricate nature (Fahrig et al., 2011; Martin et al., 2019). Thus, we used ortho-photos and *in situ* visits to select farms located in a wide range of landscape complexity and classified *a priori* the 20 localities into the three categories of landscape, frequently used to test landscape-related hypotheses (e.g. Concepción et al., 2012), namely: simple, intermediate and complex. Afterwards, we used the most recent land use cartography (SIOSE, http://www.siose.es) and the FRAGSTATS software (McGarigal et al., 2012) to calculate, in each locality, 12 metrics of landscape compositional (patch richness, diversity and evenness, percentage of semi-natural habitat cover, percentage of olive groves cover) and configurational (edge density, largest patch area, mean patch area, shape of the mean patch, Euclidean distance between nearest neighbor patches of similar uses, contagion and interspersion/juxtaposition index) heterogeneity. Then, we validated the *a priori* perceptual classification, using a classification and regression tree (CART, De’ath & Fabricius, 2000). The CART rendered that just three of these variables: percentage of semi-natural area, distance to the nearest neighbor, and mean patch area, adequately ascribed the 20 localities into the previously a priori groups (Rey et al., 2019, for more details on the procedure and thresholds). Landscape complexity metrics are available in Mendeley data (DOI: 10.17632/dchz48kfbh.1).

### Ant surveys

We sampled ant communities on 40 olive farms. At each farm, we established 12 permanent plots, four of them sited in non-crop patches (mostly remnants of semi-natural habitat) and eight inside the olive orchard matrix. In each plot, we set a pitfall trap (transparent plastic glass; 7cm of diameter and 12cm depth), filled with water, propylene glycol (1:1 proportion) and 5-10 soap drops. Every month, from April to November 2016, we collected the ants, refilled the pitfall trap and determined the collected individuals in the laboratory, under the stereomicroscope. In total, we used 3360 pitfall traps, distributed in 480 permanent plots.

### Ant species specific response

We calculated ant species specific response against the olive moth and the olive fly (i.e. appetence for these pest species) by offering real pest larvae or pupae (which were manually collected in our study areas previously the same day) to foraging ants. This experiment was conducted in two consecutive years (2018: in nine olive groves, and 2019: in 18 olive groves), during the period when these pests are more vulnerable to ants (spring for the olive moth and end of autumn for the olive fly). Thus, we offered pest larvae/pupae to actively foraging ants and recorded their response to the offerings (see Fig.1). A positive response was recorded when an ant interacted with the pest (i.e. touching it with the antennae) and bit it, taking it away (or trying to). In contrast, an ant disregarding the larvae/pupae after contacting it was considered as a negative response. Ant individuals that contacted with the pest offered and responded either positively or negatively, were collected and identified to the species level at the lab. Ant species-specific response was calculated as the proportion of positive responses (ratio positive to total offerings) against the pest species (see Garrido et al., 2002 for similar approach). Overall, 1225 offerings were recorded, 830 for the olive moth, and 395 for the olive fly.

*Tapinoma nigerrimum* alone might play an important role on pest control in olive groves (Morris et al., 2002). It was a very abundant and active species during our offerings (for both the olive moth and the olive fly), its specific response was highly positive, and was among the most abundant and widespread species in the ant assemblages of all the studied farms. Therefore, we modeled its abundance across landscape and management gradients.

We measured the ant-species specific responses to each pest species only for the species co-occurring with the vulnerable stages of the pests. Specifically, for the response against the olive moth we recorded specific responses against the anthophagous larvae for 25 out of 58 ant species sampled in the whole year (representing the 83% of the ant community abundance, including crop and non-crop plots). In this case, offerings were performed to ants patrolling olive branches and twigs. Regarding the response against the olive fly, we recorded the response of 22 out of 39 species during autumn (representing the 96% of the ant community abundance in autumn) and offerings were carried out on the ground. In some cases, we could not achieve enough offerings to obtain a robust inference of a species’ response against the pest (intraspecific variability on the response). In these cases, we decided to include these species to maximize the available information, assuming a close to binomial-type response (e.g. can predate or not; 1 positive response equals 100% positive responses). Note that very few offerings often meant a low abundance of the target species (see Table S1 in Supporting Material for the number of offerings per species and their abundance).

### Ant community predation function

Assuming a lineal increase of encounter probability and feeding needs with abundance, we expect also that predation pressure increases linearly with ant abundance (i.e. each foraging ant from the same species contributes the same to predation) and appetence for the pest by each ant species. Therefore, following Martínez-Núñez & Rey (2020a) we calculated the Ant-community Predation Function (APF) for a given pest accounting for the quantitative and qualitative components of the predation function. APF represents thus the expected predation pressure that the whole community of ants can exert over a given pest on a given farm in the considered period and, therefore, is our estimation of the ant-mediated pest control service.

We calculated the ant-community predation function against each pest, using the Equation 1.

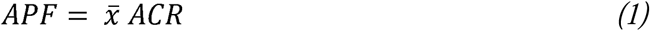

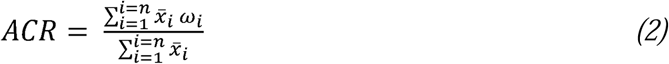

Thus, the ant predation function equals the mean abundance per pitfall of the ant community 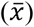 times the Ant Community Response (ACR), which is the expected probability of attack by a randomly chosen individual (last term in Eq, 1). The ACR (Eq. 2) is the sum (from species 1 to *n*) of the mean abundance of each ant species with one or more offerings, multiplied by its specific response to the pest (*ω*_i_) and divided by the total mean abundance of species with offerings. ACR as estimated here is meaningful since contains information for each ant assemblage of the *qualitative component* of the depredation function, based on its species composition and the species-specific response (Garrido et al., 2002) to pests, and on the *quantitative component* of such function (Martínez-Núñez & Rey, 2020a) based on the overall ant abundance in such community (as determined by the mean number of ants collected per pitfall in each farm).

### Artificial prey experiment

Dummy plasticine models have frequently been used to mimic potential prey, and compare experimentally attack rates by natural enemies across environmental gradients (e.g. Milligan et al., 2016). Here, we used models previously macerated in pest larvae juice. We previously recorded that ants showed a higher preference towards these dummies compared to non-macerated dummies, and also confirmed the absence of different responses against the used models and true larvae. Then, we built small exclosures to enable ant attacks, precluding removal of the plasticine larvae from other predators (see Fig. 1). We built the exclosures using small plastic Petri plates of 5cm diameter and 2cm deep. We opened two gaps (“doors”) in the plates to let ants enter and exit. Finally, a lid did protect the larvae from the rain and other animals. We set two dummies per plate, and 20 plates per farm (ten in trees adjacent to semi-natural patches, and ten in trees within the olive tree matrix, approximately 200 m distant from these patches). We conducted this experiment during two consecutive years (2018-2019), for the two pest species. In 2018, we ran this experiment in 9 farms (720 plasticine models, 360 mimicking each pest species), increasing this number to 18 farms (1440 plasticine models, 740 for each pest) in 2019. Overall, we provided and checked 2160 plasticine models. We set the experiment for the olive moth in May, targeting the anthophagous larvae (the most vulnerable to ant predation). Accordingly, the plates with the dummies were set on branches (see Fig. 1). For the olive fly, we set the experiment in November, and the plates were located on the ground, attached with metal hooks.

**Figure 1:**
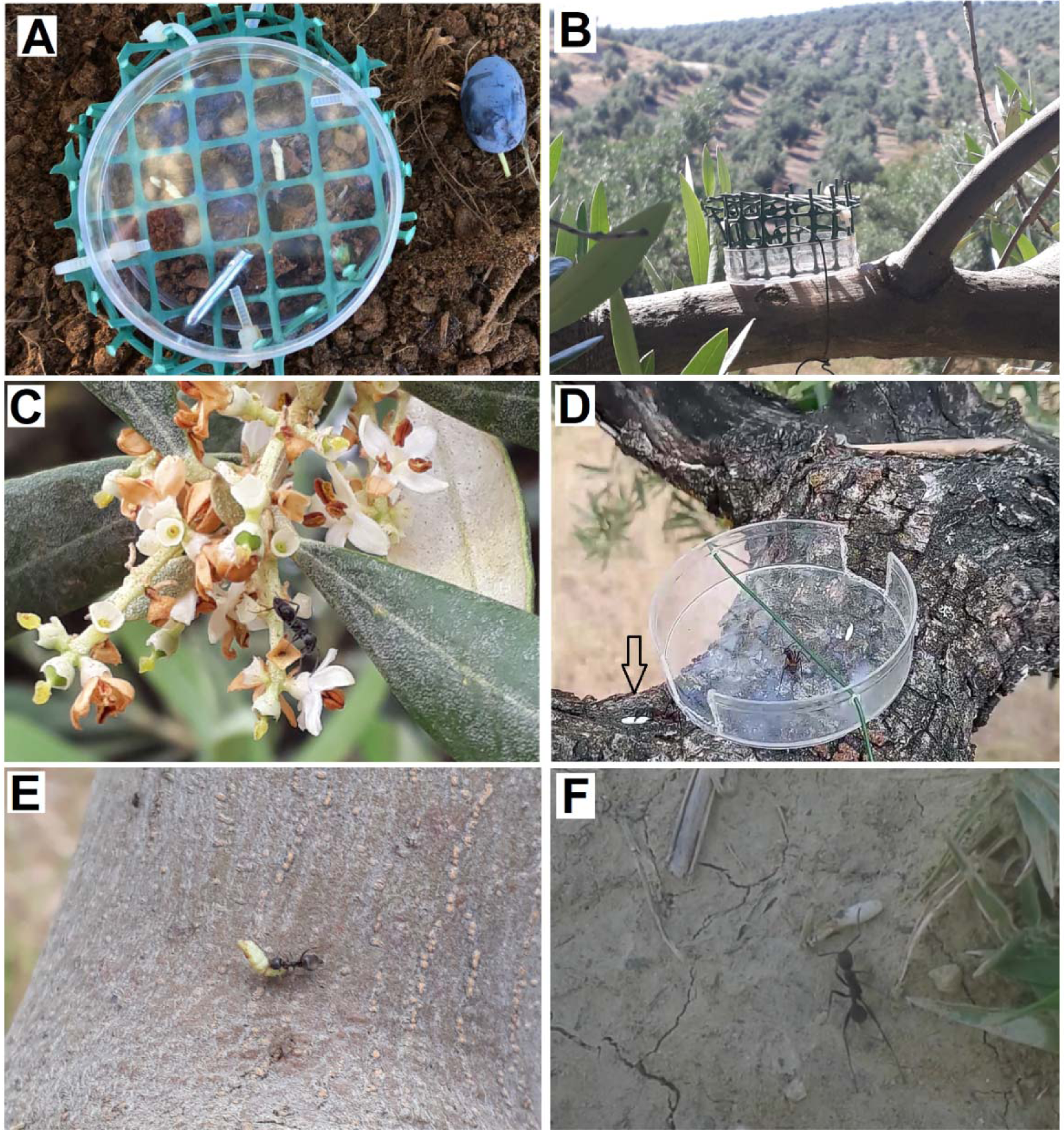
Pictures showing the experiments conducted to assess ant specific response and attack rates to plasticine dummies. A) Exclusion with plasticine models to assess attack rates by ants to the olive fly larvae, picture from above. B) Exclusion with plasticine models to assess attack rates by ants to the olive moth anthophagous larvae. C) Ant actively foraging on olive tree flowers. D) Ant predating and taking a plasticine dummy from the exclusion cage (note that the lid is missing because the cage was not completely set yet). E) Positive response to in situ offering. F) Ant contacting a larva offered, right before discovering whether it has a positive or negative response.

In this study, we assume that a higher potential predation pressure (i.e. a higher abundance of ants with a higher appetence for the pests) will lead to a higher potential pest control and less crop damage.

### Statistical analyses

First, we explored variations in ant abundance, *T. nigerrimum* abundance, and APF across farms with different management (intensive vs. extensive) and landscape complexity contexts (simple, intermediate, complex). We considered both the whole sampling period (pitfall trap averaged for the 7 months sampling period), when some stages of the olive moth can be predated by ants, and autumn (pitfall trap averaged for October and November), when the olive fly larvae are vulnerable to ants from the soil. We used the package *lme4* (Bates, Mächler, Bolker, & Walker, 2015) in R (R Core Team 2019) to run three general linear mixed models for each response variable (ant abundance, *T. nigerrimum* abundance and APF), with landscape complexity, agricultural management, and their interaction as fixed factors. We did not include ant richness in this test because it was covered elsewhere (Rey et al. 2019).

Second, we tested the effect of ecological contrast on ant abundance, ant richness, *T. nigerrimum* abundance and APF. For this, we fitted separate models for each response variable, introducing management (intensive vs. extensive) and ecological contrast (low vs. high) as an interaction fixed term.

For all pest control-related analyses, we only considered the eight plots of each farm located within the crop, to avoid confounding factors (i.e. plots not exposed to management filtering, such as the non-crop plots). However, we also tested differences between crop and non-crop plots across our management and landscape gradient to unveil possible variations in habitat preferences (see Supporting information).

All models (unless previously specified), included a locality and farm ID random factor. Model assumptions were checked by inspection of the residuals. Response variables were transformed when needed to conform to model assumptions. Results were interpreted by comparing the AIC with a null model (a model without any fixed factor), acknowledging the marginal R^2^, and post hoc Tukey multiple comparison tests with alpha = 0.10, to also account for marginally significant differences.

## Results

In total, 166611 ant individuals were collected and determined on the 40 farms (ranging between 1284 and 15775 individuals per farm). Overall, 58 different ant species, belonging to 18 different genera, were detected (ranging from 16 to 33 species per farm). A complete list of the species can be found in Table S1. *T. nigerrimum* (15.8%), *Aphaenogaster senilis* (12.5%) and *Messor barbarus* (12.1%) were the most abundant and ubiquitous species. They occurred on 36, 37 and 38 out of 40 farms respectively).

The ant-species specific response varied between 13% positive response from *Pheidole pallidula* or 16% from *Plagiolepis schitzii*, and 100% positive response from *Camponotus barbaricus* for olive moth. Against the olive fly, it varied between 0% from *Plagiolepis pygmaea* and 100% from *Aphaenogaster iberica* (Table S1). Some species were very abundant/active during the experiments and had a clear positive response against both pest species, for instance, *Aphaenogaster senilis, Tapinoma nigerrimum* and some species belonging to the genera *Camponotus* or *Lassius*. Smaller species, belonging to the genera *Plagiolepis, Pheidole* or *Tetramorium* had very few positive responses (Table S1). The *T. nigerrimun* response to offerings was among the highest, 90% for the olive moth and 93% for the olive fly.

### Landscape and management effects on ants

Mean ant abundance per pitfall/farm varied between 19 and 149 individuals per trap, and was higher on farms with extensive management of intermediate landscapes (see Table 1 and Fig. 2). Differences across farms with contrasting management were also stronger at intermediate landscapes. The mean abundance of *T. nigerrimum* per pitfall/farm during the whole study period varied between 0 and 40 and was not affected by management or landscape (see Table 1 and Fig. 2). APF varied between 14 in Olivar de la Luna intensive, and 97 in Virgen de los Milagros extensive for the olive moth and between 11 in Olivar de la Luna intensive and 99 in Los Ojuelos extensive, for the olive fly. The APF for the olive moth was affected by management and was on average a 30% higher on farms under extensive management (see Table 1 and Fig. 2).

**Table 1:** Linear mixed models showing the importance of Management, Landscape and Management*Landscape interaction to explain variations in Ant abundance, *T. nigerrimum* abundance and APF with data for the whole year (olive moth) and only October-November (olive fly). In bold, significant models, with alpha < 0.05. In italics, marginally significant models, with 0.05 < alpha < 0.1.

| Response<br>variable | Explanatory variable | $\Delta AIC$<br>NULL | $X^2$ | P | Marginal<br>$R^2$ | Conditional<br>$R^2$ |
| --- | --- | --- | --- | --- | --- | --- |
| Ant abundance<br>(moth) | **Management** | **4.12** | **6.16** | **0.013** | **4.0** | **27.5** |
|  | *Landscape* | *1.43* | *5.43* | *0.066* | *4.7* | *27.8* |
|  | **Management\*Landscape** | **5.3** | **15.30** | **0.009** | **10.2** | **28.9** |
| T. nigerrimum<br>abundance<br>(moth) | Management | -1.7 | 0.21 | 0.648 | 0.1 | 32.0 |
|  | Landscape | 0.0 | 3.92 | 0.141 | 4.6 | 33.2 |
|  | Management*Landscape | -2.7 | 7.29 | 0.200 | 6.4 | 34.4 |
| APF (moth) | **Management** | **2.8** | **4.79** | **0.029** | **2.9** | **22.9** |
|  | Landscape | -0.2 | 3.78 | 0.151 | 3.0 | 23.6 |
|  | *Management\*Landscape* | *0.3* | *10.3* | *0.067* | *6.6* | *24.9* |
| Ant abundance<br>(fly) | Management | -0.67 | 1.33 | 0.248 | 0.8 | 23.6 |
|  | Landscape | -0.10 | 3.89 | 0.143 | 4.1 | 24.9 |
|  | Management*Landscape | -4.52 | 5.48 | 0.360 | 4.9 | 26.1 |
| T. nigerrimum<br>abundance (fly) | Management | -1.92 | 0.08 | 0.779 | 0.1 | 43.3 |
|  | *Landscape* | *1.63* | *5.63* | *0.059* | *7.9* | *44.0* |
|  | Management*Landscape | -2.84 | 7.16 | 0.209 | 9.1 | 46.2 |
| APF (fly) | Management | -1.6 | 0.39 | 0.530 | 0.3 | 28.6 |
|  | *Landscape* | *0.8* | *4.8* | *0.091* | *7.3* | *29.4* |
|  | Management*Landscape | -3.33 | 6.6 | 0.247 | 8.5 | 30.2 |

**Figure 2:**
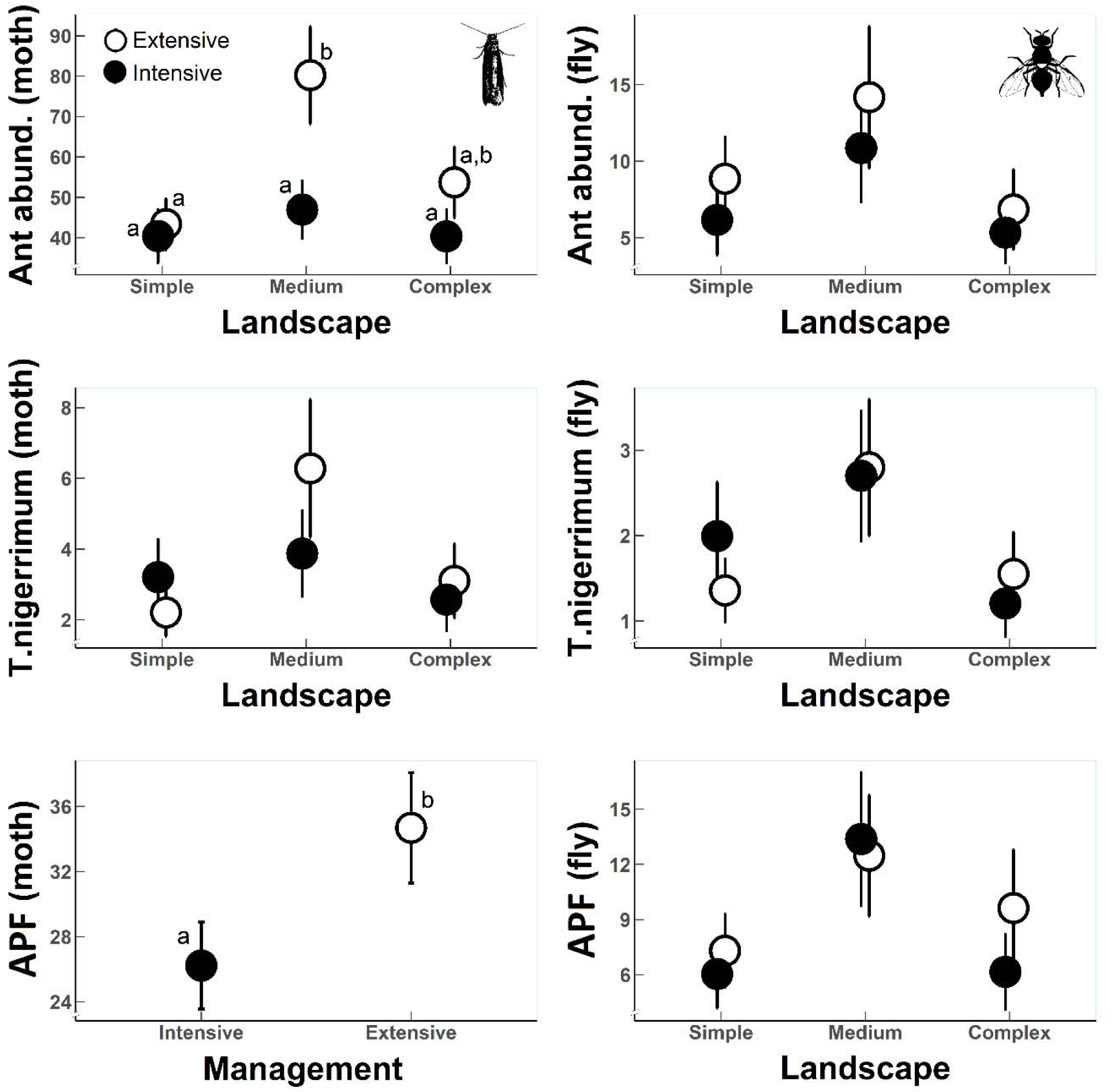
Landscape and management effects on ant abundance, *T. nigerrimum* abundance and APF for the olive moth (all sampling period) and the olive fly (only data from October and November). Only the best model is represented, based on AIC. In the case that no model was better than the null model, the model with the highest marginal R^2^ was plotted. Letters show Tukey post hoc multiple comparison test with an alpha=0.1 when statistically significant differences exist.

Across farms, *T. nigerrimum* accounted for 17±2 % (Mean±1SE) of the APF against the olive moth (range 0-50% across farms). During the months of October and November (when the olive fly is vulnerable to ants), *T. nigerrimum* accounted for 8±3 % of the APF against the olive fly (range 0-67%). No remarkable differences were detected in Autumn for abundance, APF on the moth or APF on the fly across our management and landscape complexity categories. However, marginally significant effects of landscape were detected for *T. nigerrimum* abundance and APF, peaking at intermediate landscapes.

Mean differences in species abundance, species richness, *T. nigerrimum* abundance and APF (annual) tended to be greater under high ecological contrast, however, these differences were only remarkable for ant abundance and ant richness (see Fig. S1 and Table S2 in Supporting Information). *T. nigerrimum* was not significantly affected by ecological contrast and APF showed consistently higher values in extensive management, although differences were slightly stronger between pairs with high contrast (Fig. S1; effect size of 1.25 in low contrast vs. 1.38 in high contrast).

### Ant attack rates

Attack rates to plasticine models averaged 27% (ranging between 10% and 70%) for the olive moth, and 20% (ranging between 0% and 67% for the fly). Attack rates to the olive moth models were higher on farms with extensive management (see Table 2 and Figure 3). However, intensity of the ecological contrast by different management did not drive important differences on attack rates for the olive fly models. Closeness to semi-natural habitat did not explain differences in attack rates under extensive/intensive farming and for any pest species (see Table 2).

**Table 2:** Linear mixed models showing the importance of Management, Distance to patches of SNH, and their interaction to explain variations in observed attack rates to the moth and the fly. In bold, significant models, with alpha = 0.05. In italic, marginally significant models, with alpha = 0.1.

| **Response**<br>**variable** | **Explanatory variable** | **ΔAIC**<br>**NULL** | **$X^2$** | **P** | **Marginal**<br>**$R^2$** | **Conditional**<br>**$R^2$** |
| --- | --- | --- | --- | --- | --- | --- |
| Attack rates<br>(moth) | **Management** | **4.44** | **6.44** | **0.011** | **5.8** | **17.2** |
|  | Distance to SNH | -0.75 | 1.26 | 0.262 | 0.3 | 17.2 |
|  | *Management\*Distance* | *1.73* | *7.73* | *0.052* | *6.2* | *17.5* |
| Attack rates (fly) | Management | 0.21 | 2.21 | 0.137 | 2.0 | 21.4 |
|  | Distance to SNH | -1.81 | 0.19 | 0.662 | 0.0 | 21.3 |
|  | Management*Distance | -2.05 | 3.00 | 0.268 | 2.2 | 19.2 |

**Figure 3:**
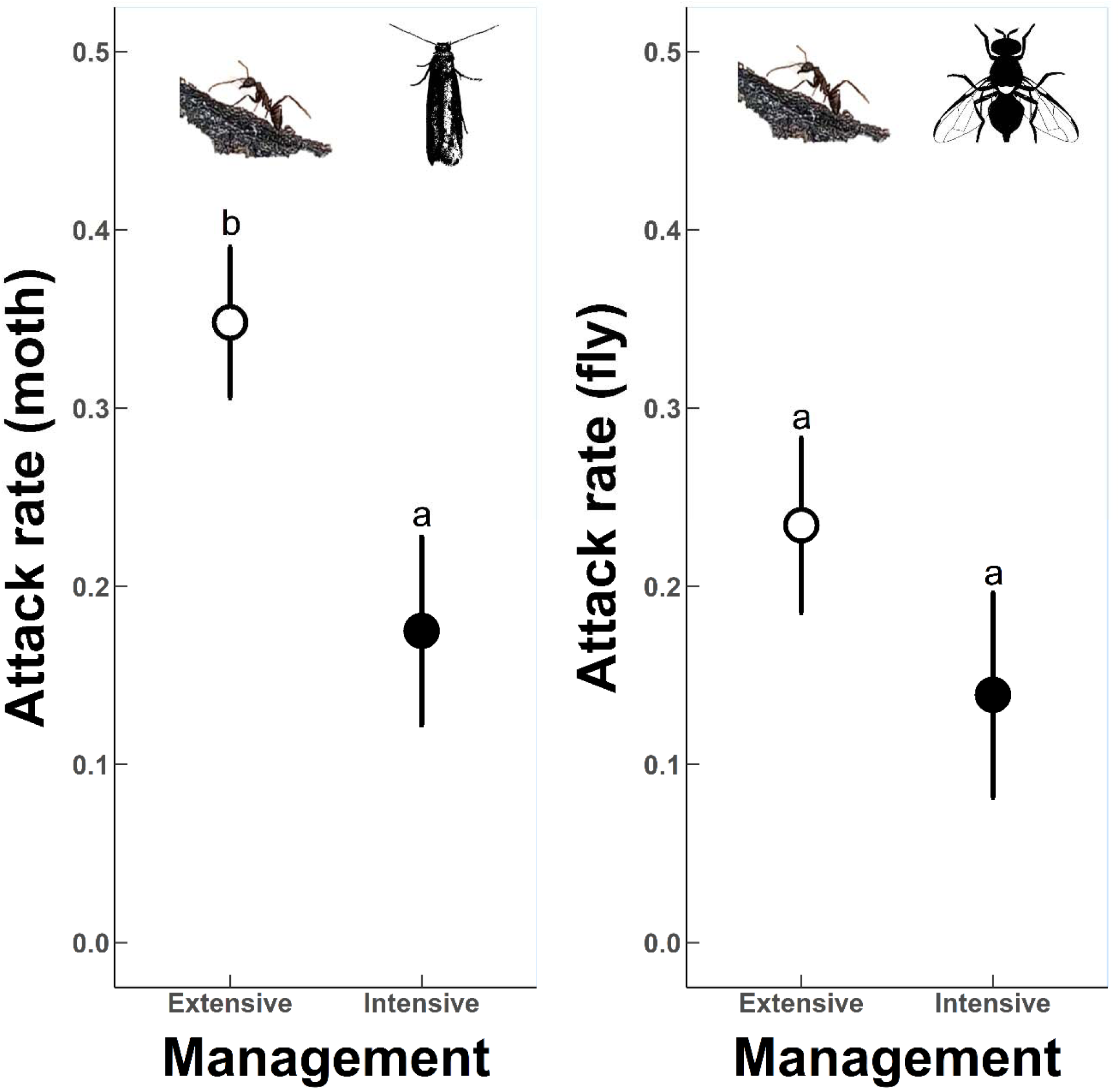
Management effects on attack rates to plasticine dummies mimicking olive moth (left) and olive fly (right) larvae. Letters show Tukey post hoc multiple comparison test with an alpha=0.1.

## Discussion

Unlike permanent agroecosystems in tropics, drivers of ant-mediated pest control have received little attention in dryland permanent agroecosystems, that cover extensive areas (http://www.fao.org/faostat). Using as study system Mediterranean olive groves, the most extended arboreal cropland in drylands, we quantitatively report, for the first time, that ants have a strong potential for pest control in this agroecosystem at large regional scale. We predicted interacting effects of agricultural management and landscape heterogeneity on ant community and ant-mediated predation pressure. Our results provided general support for our predictions. Ant abundance was higher on farms with extensive management and intermediate landscape complexity, but did not increase linearly with landscape complexity. Also, the ant-community predation function (APF) was higher on extensively managed farms. However, these patterns were not observed in autumn, when activity of ants naturally decays. Accordingly, only for the olive moth a higher predation pressure was patent in extensive farms, as shown by higher attack rates to plasticine models. Interestingly, differences in APF matched differences on attack rates to plasticine models. We further confirmed that the increase in the intensity of the ecological contrast generated by agri-enviromental measures rendered remarkable increase in ant abundance and ant richness, although such differences barely transferred to the predation function. Last, *T. nigerrimum* was one of the most important contributors to the pest predation function and its abundance was not evidently affected by management nor landscape complexity, as it could be expected from its tolerance to disturbance (see Table S1; Redolfi et al.,1999).

### Management effects on ant community and APF

The overall positive effect of herb cover extensification on ant abundance and potential predation pressure (i.e. APF) here detected confirms the positive relation between local habitat heterogeneity and ant diversity previously reported in other agrosytems. For instance, ant diversity diminished with cacao and coffee intensification (Philpott & Armbrecht, 2006), with herb cover intensification in olive groves (Rey et al., 2019), and distance to natural vegetation strips in sugarcane (Rivera-Pedroza et al., 2019). The presence of plant cover, thus, seems to be important for ants because it enhances habitat suitability. Herb cover might benefit ant communities in different ways, such as, buffering extreme temperatures in summer and winter, increasing the presence of alternative prey or providing direct feeding resources via floral and extrafloral nectaries (Jones, Koptur, & von Wettberg, 2017). Frequent soil disturbances in intensive farms, due to tillage or erosion by rain, might drive part of the observed effects as well. Other studies found negative effect of tillage on ants (Sharley, Hoffmann, & Thomson, 2008; but see Rodríguez, González, & Campos, 2012). Also, ant activity is considerably reduced due to phenology and climatic conditions in autumn, possibly homogenizing the communities of active ants in extensive and intensive farms. However, other natural enemies might be important, and some authors have demonstrated how soil disturbance in autumn leads to reduced olive fly predation by soil arthropods (Ortega et al., 2018). As other studies have previously suggested (Morris et al., 2002), *T. nigerrimum* appears in our study as a very abundant and ubiquitous species in olive groves at the regional scale. Moreover, the experiments showed a high frequency of this species during the key periods (when pests are in more vulnerable stages: spring on the trees, autumn on the ground), with an active foraging behavior, and a high appetence for both pest species (the olive moth and the olive fly). Therefore, this ant species is likely an important natural enemy for the olive moth and the olive fly, which presence and abundance is desirable in olive groves. As previously reported by other authors, *T. nigerrimum*, seems relatively tolerant to intensive management and landscape simplification (Redolfi et al.,1999), being a generalist and opportunistic species (Table S1; Roig & Espadaler, 2010) and might be a key species for pest control in intensive olive groves, since is the species with the overall highest contribution to APF in our study (Table S1 in Supporting Information).

### Landscape effects on ants and APF

As predicted, landscape complexity, strongly moderated (sensu Tscharstke et al. 2012) the management effects on ant abundance, and to less extent on richness and APF. Aristizábal & Metzger (2019) also showed that landscape structure can modulate ant-pest interactions, varying the pest control potential. According to the intermediate-landscape complexity hypothesis (Tscharntke et al. 2012), in our study, extensive management increased ant abundance only in intermediate landscapes. This might be explained by a combination of spillover from semi-natural habitats to the crop, and differential use preference of crop and non-crop habitats in different landscape contexts. However, differences in ant abundance between crop and non-crop plots did not vary across landscapes (although richness did, see Fig. S2 in Supporting information), indicating that this pattern is not a matter of habitat preference. We argue that intermediate landscapes provide intermediate disturbance/heterogeneity, favoring non-dominance and coexistence of generalist and tolerant species at higher densities. Other authors have shown that habitat and landscape complexity affect natural enemy abundance (Álvarez et al., 2019) and overall predation pressure by soil arthropods on olive pests (Ortega et al., 2018) which suggests that additive effects might be important.

### Ecological contrast effects on ants and APF

Ant abundance and richness were negatively affected also by the intensity of the ecological contrast, showing that in the olive groves from Andalusia ants are affected by the use of pesticides or synthetic fertilizers. Likewise, a study in Portuguese olive groves, identified *Formicidae* as one of the most sensitive taxa to insecticide application (Santos, Pereira, Torres, & Nogueira, 2007). Surprisingly, although, the intensity of the ecological contrast affected to ant abundance and richness it was not a major determinant of the ant’s depredation function on olive pests in our farms, which may be related to the fact that the most abundant and important species for pest control are generalists (e.g. *T. nigerrimum, A. senilis* or *C. barbaricus*). In any case, this result does not provide much support to the ecological contrast hypothesis for ant-mediated predation pressure in our system, hypothesis that was, however, validated on the same olive groves for solitary bees (Martínez□Núñez et al., 2020), and has found support also in other agroecosystems for pollinators (Marja et al., 2019). We suggest that contrasting results between insect pollinator and ant-mediated services in the response to the intensity of ecological contrast by AES in olive groves might be due to the more direct exposition of pollinators to pesticides compared to ground-dwelling ants.

While ant-community predation function (APF) against the olive moth increased clearly with extensive farming, the APF for the olive fly did not. Remarkably, this result matches the outcomes found for attack rate patterns to plasticine models. Attack rates were higher in extensive farms for the olive moth, but this pattern was diluted for the olive fly. This result shows that our index (i.e. the ant community response), is a fine measure of ants’ predation pressure over these two pest species, and a good predictive measure of pest control potential in olive groves. Nonetheless, there is need to acknowledge that ants, as typical generalist predators, can provide disservices as well, by predating on other potential natural enemies, or even protecting some pests from enemies (e.g. the vine mealybug in table-grave vineyards (Beltrà et al., 2017). Therefore, further research is needed in olive groves to understand their net effect when accounting for multispecies and overall pest control.

Overall, this study shows how extensive herb cover management and certain levels of landscape complexity are needed to maximize ant-driven predation pressure over pests in olive groves.

## Acknowledgements

We thank the owners of the studied olive groves. We also thank Gemma Calvo, M^a^ José Navarro, Rubén Tarifa, José L. Molina and Enrique Muñoz for field assistance and Francisco Valera, Carlos Ruiz and José Eugenio Gutiérrez for logistic support. This work is part of projects AGRABIES (CGL2015□68963□C2, MINECO, Gobierno de España and FEDER) and OLIVARES VIVOS (LIFE14 NAT/ES/001094, European Commission). CMN was granted a predoctoral fellowship (BES-2016-078736). Authors have no conflict of interests.

## Authors Contributions

CMN and PJR conceived the ideas and design of the study and led the writing of the MS. CMN, JI, AJM, FMC and PJR conducted fieldwork. TS and FMC determined ants in the lab with AJM supervision. TS processed GIS data and characterized the landscapes. CMN analyzed the data. All authors revised the final version and gave their approval for submission.

## Data availability

After acceptance, data will be freely available in a public repository.

## References

Álvarez, H. A., Morente, M., Oi, F. S., Rodríguez, E., Campos, M., & Ruano, F. (2019). Semi-natural habitat complexity affects abundance and movement of natural enemies in organic olive orchards. Agriculture, Ecosystems & Environment, 285, 106618. https://doi.org/10.1016/J.AGEE.2019.106618

Aristizábal, N., & Metzger, J. P. (2019). Landscape structure regulates pest control provided by ants in sun coffee farms. Journal of Applied Ecology, 56(1), 21–30. https://doi.org/10.1111/1365-2664.13283

Bates, D., Mächler, M., Bolker, B., & Walker, S. (2015). Fitting Linear Mixed-Effects Models Using lme4. Journal of Statistical Software, 67(1), 1–48. https://doi.org/10.18637/jss.v067.i01

Beltrà, A., Navarro-Campos, C., Calabuig, A., Estopà, L., Wäckers, F. L., Pekas, A., & Soto, A. (2017). Association between ants (Hymenoptera: Formicidae) and the vine mealybug (Hemiptera: Pseudococcidae) in table-grape vineyards in Eastern Spain. Pest Management Science, 73(12), 2473–2480. https://doi.org/10.1002/ps.4640

Bommarco, R., Kleijn, D., & Potts, S. G. (2013). Ecological intensification: Harnessing ecosystem services for food security. Trends in Ecology and Evolution. https://doi.org/10.1016/j.tree.2012.10.012

Concepción, E. D., Díaz, M., Kleijn, D., Báldi, A., Batáry, P., Clough, Y., … Verhulst, J. (2012). Interactive effects of landscape context constrain the effectiveness of local agri-environmental management. Journal of Applied Ecology, 49(3), no-no. https://doi.org/10.1111/j.1365-2664.2012.02131.x

Cotes, B., Campos, M., Pascual, F., García, P. A., & Ruano, F. (2010). Comparing taxonomic levels of epigeal insects under different farming systems in Andalusian olive agroecosystems. Applied Soil Ecology, 44(3), 228–236. https://doi.org/10.1016/j.apsoil.2009.12.011

Daane, K. M., & Johnson, M. W. (2010). Olive Fruit Fly: Managing an Ancient Pest in Modern Times. Annual Review of Entomology, 55(1), 151–169. https://doi.org/10.1146/annurev.ento.54.110807.090553

De’ath, G., & Fabricius, K. E. (2000). Classification and regression trees: a powerful yet simple technique for ecological data analysis. Ecology, 81(11), 3178–3192. https://doi.org/10.1890/0012-9658(2000)081[3178:CARTAP]2.0.CO;2

Escobar-Ramírez, S., Grass, I., Armbrecht, I., & Tscharntke, T. (2019). Biological control of the coffee berry borer: Main natural enemies, control success, and landscape influence. Biological Control, 136, 103992. https://doi.org/10.1016/J.BIOCONTROL.2019.05.011

Fahrig, L., Baudry, J., Brotons, L., Burel, F. G., Crist, T. O., Fuller, R. J., … Martin, J. L. (2011). Functional landscape heterogeneity and animal biodiversity in agricultural landscapes. Ecology Letters, 14(2), 101–112. https://doi.org/10.1111/j.1461-0248.2010.01559.x

Gallé, R., Happe, A., Baillod, A. B., Tscharntke, T., & Batáry, P. (2019). Landscape configuration, organic management, and within□field position drive functional diversity of spiders and carabids. Journal of Applied Ecology, 56(1), 63–72. https://doi.org/10.1111/jpe.2019.56.issue-1

Garrido, J. L., Rey, P. J., Cerdá, X., & Herrera, C. M. (2002). Geographical variation in diaspore traits of an ant□dispersed plant (*Helleborus foetidus*): are ant community composition and diaspore traits correlated? Journal of Ecology, 90(3), 446–455. https://doi.org/10.1046/j.1365-2745.2002.00675.x

Gkisakis, V., Volakakis, N., Kollaros, D., Bàrberi, P., & Kabourakis, E. M. (2016). Soil arthropod community in the olive agroecosystem: Determined by environment and farming practices in different management systems and agroecological zones. Agriculture, Ecosystems and Environment, 218, 178–189. https://doi.org/10.1016/j.agee.2015.11.026

Gonçalves, M. F., & Pereira, J. A. (2012). Abundance and Diversity of Soil Arthropods in the Olive Grove Ecosystem. Journal of Insect Science, 12(20), 1–14. https://doi.org/10.1673/031.012.2001

Gurr, G. M., Wratten, S. D., Landis, D. A., & You, M. (2017). Habitat Management to Suppress Pest Populations: Progress and Prospects. Annual Review of Entomology, 62(1), 91–109. https://doi.org/10.1146/annurev-ento-031616-035050

Haan, N. L., Zhang, Y., & Landis, D. A. (2019). Predicting Landscape Configuration Effects on Agricultural Pest Suppression. Trends in Ecology and Evolution. Elsevier Ltd. https://doi.org/10.1016/j.tree.2019.10.003

Jones, I. M., Koptur, S., & von Wettberg, E. J. (2017). The use of extrafloral nectar in pest management: overcoming context dependence. (S. Diamond, Ed.), Journal of Applied Ecology. Blackwell Publishing Ltd. https://doi.org/10.1111/1365-2664.12778

Kleijn, D., Rundlöf, M., Scheper, J., Smith, H. G., & Tscharntke, T. (2011). Does conservation on farmland contribute to halting the biodiversity decline? Trends in Ecology & Evolution, 26(9), 474–481. https://doi.org/10.1016/J.TREE.2011.05.009

Malheiro, R., Casal, S., Baptista, P., & Pereira, J. A. (2015). A review of Bactrocera oleae (Rossi) impact in olive products: From the tree to the table. Trends in Food Science and Technology. Elsevier Ltd. https://doi.org/10.1016/j.tifs.2015.04.009

Marja, R., Kleijn, D., Tscharntke, T., Klein, A. M., Frank, T., & Batáry, P. (2019). Effectiveness of agri-environmental management on pollinators is moderated more by ecological contrast than by landscape structure or land-use intensity. (J. Knops, Ed.), Ecology Letters. Blackwell Publishing Ltd. https://doi.org/10.1111/ele.13339

Martin, E. A., Dainese, M., Clough, Y., Báldi, A., Bommarco, R., Gagic, V., … Steffan□Dewenter, I. (2019). The interplay of landscape composition and configuration: new pathways to manage functional biodiversity and agroecosystem services across Europe. Ecology Letters, 22(7). https://doi.org/10.1111/ele.13265

Martínez-Núñez, C., & Rey, P. J. (2020a). Assessing the predation function via quantitative and qualitative interaction components. BioRxiv, 2020.05.16.099721. https://doi.org/10.1101/2020.05.16.099721

Martínez□Núñez, C., Manzaneda, A. J., Isla, J., Tarifa, R., Calvo, G., Molina, J. L., … Rey, P. J. (2020b). Low□intensity management benefits solitary bees in olive groves. Journal of Applied Ecology, 57(1), 111–120. https://doi.org/10.1111/1365-2664.13511

McGarigal, K., SA Cushman, and E Ene. 2012. FRAGSTATS v4: Spatial Pattern Analysis Program for Categorical and Continuous Maps. University of Massachusetts, Amherst.

Milligan, M. C., Johnson, M. D., Garfinkel, M., Smith, C. J., & Njoroge, P. (2016). Quantifying pest control services by birds and ants in Kenyan coffee farms. Biological Conservation, 194, 58–65. https://doi.org/10.1016/J.BIOCON.2015.11.028

Morris, J., Vandermeer, J., & Perfecto, I. (2015). A keystone ant species provides robust biological control of the coffee berry borer under varying pest densities. PLoS ONE, 10(11). https://doi.org/10.1371/journal.pone.0142850

Morris, J., Jimenez-Soto, E., Philpott, S., & Perfecto, I. (2018). Ant-mediated (Hymenoptera: Formicidae) biological control of the coffee berry borer: diversity, ecological complexity, and conservation biocontrol. MYRMECOLOGICAL NEWS, 26, 1–17.

Morris, T. I., Symondson, W., Kidd, N., Jervis, M., & Campos, M. (1998). Are ants significant predators of the olive moth, Prays oleae? Crop Protection, 17(4), 365–366. https://doi.org/10.1016/S0261-2194(98)00016-7

Morris, T. I., Symondson, W. O. C., Kidd, N. A. C., & Campos, M. (2002). The effect of different ant species on the olive moth, Prays oleae (Bern.), in Spanish olive orchard. Journal of Applied Entomology, 126(5), 224–230. https://doi.org/10.1046/j.1439-0418.2002.00647.x

Offenberg, J. (2015). Ants as tools in sustainable agriculture. Journal of Applied Ecology, 52(5), 1197–1205. https://doi.org/10.1111/1365-2664.12496

Orsini, M. M., Daane, K. M., Sime, K. R., & Nelson, E. H. (2007). Mortality of olive fruit fly pupae in California. Biocontrol Science and Technology, 17(8), 797–807. https://doi.org/10.1080/09583150701527359

Ortega, M., Sánchez-Ramos, I., González-Núñez, M., & Pascual, S. (2018). Time course study of Bactrocera oleae (Diptera: Tephritidae) pupae predation in soil: the effect of landscape structure and soil condition. Agricultural and Forest Entomology, 20(2), 201–207. https://doi.org/10.1111/afe.12245

Paredes, D., Karp, D. S., Chaplin-Kramer, R., Benítez, E., & Campos, M. (2019). Natural habitat increases natural pest control in olive groves: economic implications. Journal of Pest Science, 92(3), 1111–1121. https://doi.org/10.1007/s10340-019-01104-w

Pelekassis, C. D. (1962). A contribution to the study or nomenclature, taxonomy, biology, ecology and the natural parasitisation of the olive kernel borer ()Prays oleae (Bernard) Lesne). A Contribution to the Study or Nomenclature, Taxonomy, Biology, Ecology and the Natural Parasitisation of the Olive Kernel Borer ()Prays Oleae (Bernard) Lesne).

Philpott, S. M., & Armbrecht, I. (2006). Biodiversity in tropical agroforests and the ecological role of ants and ant diversity in predatory function. Ecological Entomology, 31(4), 369–377. https://doi.org/10.1111/j.1365-2311.2006.00793.x

Picchi, M. S., Marchi, S., Albertini, A., & Petacchi, R. (2017). Organic management of olive orchards increases the predation rate of overwintering pupae of Bactrocera oleae (Diptera: Tephritidae). Biological Control, 108, 9–15. https://doi.org/10.1016/j.biocontrol.2017.02.002

R Core Team (2019). R: A language and environment for statistical computing. R Foundation for Statistical Computing, Vienna, Austria. URL http://www.R-project.org/.

Redolfi, I., Tinaut, A., Pascual, F., & Campos, M. (1999). Qualitative aspects of myrmecocenosis (Hym., Formicidae) in olive orchards with different agricultural management in Spain. Journal of Applied Entomology, 123(10), 621–627. https://doi.org/10.1046/j.1439-0418.1999.00411.x

Rey, P. J., Manzaneda, A. J., Valera, F., Alcántara, J. M., Tarifa, R., Isla, J., … Ruiz, C. (2019). Landscape-moderated biodiversity effects of ground herb cover in olive groves: Implications for regional biodiversity conservation. Agriculture, Ecosystems & Environment, 277, 61–73. https://doi.org/10.1016/J.AGEE.2019.03.007

Rivera-Pedroza, L. F., Escobar, F., Philpott, S. M., & Armbrecht, I. (2019). The role of natural vegetation strips in sugarcane monocultures: Ant and bird functional diversity responses. Agriculture, Ecosystems and Environment, 284, 106603. https://doi.org/10.1016/j.agee.2019.106603

Rodríguez, E., González, B., & Campos, M. (2012). Natural enemies associated with cereal cover crops in olive groves. Bulletin of Insectology, 65(1), 43–49.

Roig & Spadaler (2010). Proposal of functional groups of ants for the Iberian Peninsula and Balearic Islands, and their use as bioindicators. Iberomyrmex, 2, 2010.

Ruano, F., Lozano, C., Garcia, P., Peña, A., Tinaut, A., Pascual, F., & Campos, M. (2004). Use of arthropods for the evaluation of the olive-orchard management regimes. Agricultural and Forest Entomology, 6(2), 111–120. https://doi.org/10.1111/j.1461-9555.2004.00210.x

Santos, S. A. P., Cabanas, J. E., & Pereira, J. A. (2007). Abundance and diversity of soil arthropods in olive grove ecosystem (Portugal): Effect of pitfall trap type. European Journal of Soil Biology, 43(2), 77–83. https://doi.org/10.1016/j.ejsobi.2006.10.001

Santos, S. A. P., Pereira, J. A., Torres, L. M., & Nogueira, A. J. A. (2007). Evaluation of the effects, on canopy arthropods, of two agricultural management systems to control pests in olive groves from north-east of Portugal. Chemosphere, 67(1), 131–139. https://doi.org/10.1016/j.chemosphere.2006.09.014

Scheper, J., Holzschuh, A., Kuussaari, M., Potts, S. G., Rundlöf, M., Smith, H. G., & Kleijn, D. (2013). Environmental factors driving the effectiveness of European agri-environmental measures in mitigating pollinator loss - a meta-analysis. Ecology Letters, 16(7), 912–920. https://doi.org/10.1111/ele.12128

Sharley, D. J., Hoffmann, A. A., & Thomson, L. J. (2008). The effects of soil tillage on beneficial invertebrates within the vineyard. Agricultural and Forest Entomology, 10(3), 233–243. https://doi.org/10.1111/j.1461-9563.2008.00376.x

Thurman, J. H., Northfield, T. D., & Snyder, W. E. (2019). Weaver Ants Provide Ecosystem Services to Tropical Tree Crops. Frontiers in Ecology and Evolution, 7, 120. https://doi.org/10.3389/fevo.2019.00120

Tscharntke, T., Tylianakis, J. M., Rand, T. A., Didham, R. K., Fahrig, L., Batáry, P., … Westphal, C. (2012). Landscape moderation of biodiversity patterns and processes - eight hypotheses. Biological Reviews, 87(3), 661–685. https://doi.org/10.1111/j.1469-185X.2011.00216.x

Way, M. J., & Khoo, K. C. (1992). Role of Ants in Pest Management. Annual Review of Entomology, 37(1), 479–503. https://doi.org/10.1146/annurev.en.37.010192.002403

