## Supporting Material for "Ant community potential for pest control in olive groves: management and landscape effects"

**Journal of Applied Ecology**

^1^ Dept. Biología Animal, Biología Vegetal y Ecología, Universidad de Jaén. E-23071 Jaén, Spain.

^2^ Instituto Interuniversitario del Sistema Tierra de Andalucía, Universidad de Jaén, E-23071 Jaén, Spain.

^3^ Centro de Estudios Avanzados en Ciencias de la Tierra, Energía y Medio Ambiente. Universidad de Jaén, E-23071 Jaén, Spain.

^4^ Integrative Ecology Group, Estación Biológica de Doñana, EBD-CSIC, Sevilla, Spain.

Declaration of interest: None

**Table S1:** Ant species, their abundance, number of positive responses, number of total offerings and their dietary/habitat preferences. Overall contribution to APF (for prays) and functional groups according to Roig and Espadaler (2010). Code corresponds to: Criptic (C), Generalist and/or Opportunist (GO), Hot Climate Specialist and/or Open Habitat (HCS/OH), Cold Climate Specialist and/or Shadow Habitat (CCS/SH) and Dead Wood Specialist (CWDS).

| **Ant Species** | **Abundance** | **Positive responses (offerings) against the moth** | **Positive responses (offerings) against the fly** | **Overall contribution to APF (all farms together)** | **Ant Functional Group** |
| --- | --- | --- | --- | --- | --- |
| Aphaenogaster dulcinaea | 11 | -- | -- | -- | C |
| Aphaenogaster gemella | 52 | -- | -- | -- | GO |
| Aphaenogaster gibbosa | 300 | -- | -- | -- | C |
| Aphaenogaster ibérica | 2672 | 0.96 (51) | 1 (14) | 2567 | GO |
| Aphaenogaster senilis | 20851 | 0.90 (71) | 0.84 (62) | 18795 | GO |
| Aphaenogaster splendida | 557 | -- | -- | -- | C |
| Aphaenogaster striativentris | 244 | -- | -- | -- | HCS/OH |
| Aphaenogaster subterránea | 2776 | -- | -- | -- | C |
| Camponotus aethiops | 786 | 0.79 (75) | 0.5 (4) | 618 | HCS/OH |
| Camponotus barbaricus | 4 | 1 (22) | 0 (2) | 4 | HCS/OH |
| Camponotus cruentatus | 560 | -- | 1 (1) | -- | HCS/OH |
| Camponotus figaro | 16 | 1 (1) | -- | 16 | HCS/OH |
| Camponotus foreli | 989 | 0.63 (59) | 1 (1) | 620 | HCS/OH |
| Camponotus lateralis | 67 | 1 (3) | -- | 67 | CCS/SH |
| Camponotus ligniperdus | 3 | -- | -- | -- | CWDS |
| Camponotus piceus | 14 | -- | 1 (2) | -- | HCS/OH |
| Camponotus pilicornis | 65 | -- | -- | -- | HCS/OH |
| Camponotus sylvaticus | 16282 | 0.86 (73) | 1 (2) | 14052 | HCS/OH |
| Cardiocondyla batesii | 138 | -- | -- | -- | GO |
| Cardiocondyla mauritanica | 6 | -- | -- | -- | GO |
| Cataglyphis hispánica | 329 | 1 (2) | -- | -- | HCS/OH |
| Cataglyphis ibérica | 67 | -- | -- | -- | HCS/OH |
| Cataglyphis rosenhaueri | 12911 | 1 (3) | 1 (1) | 12911 | HCS/OH |
| Cataglyphis velox | 2480 | 1 (2) | 1 (1) | 2480 | HCS/OH |
| Crematogaster auberti | 1495 | 0.92 (51) | 0.14 (21) | 1378 | GO |
| Crematogaster scutellaris | 832 | 1 (12) | -- | 832 | GO |
| Crematogaster sordidula | 429 | -- | -- | -- | GO |
| Formica cunicularia | 287 | -- | -- | -- | HCS/OH |
| Formica gerardi | 27 | -- | -- | -- | HCS/OH |
| Formica subrufa | 8351 | 0.8 (20) | 0.8 (25) | 6681 | HCS/OH |
| Goniomma sp | 20 | -- | -- | -- | HCS/OH |
| Lasius brunneus | 143 | 1 (6) | -- | 143 | CCS/SH |
| Lasius flavus | 1 | -- | -- | -- | C |
| Lasius grandis | 1687 | 0.87 (30) | 1 (13) | 1462 | CCS/SH |
| Lasius lasioides | 7 | -- | -- | -- | CCS/SH |
| Lasius niger | 215 | 0.82 (28) | -- | 177 | CCS/SH |
| Messor barbarus | 20119 | -- | 0.62 (79) | -- | HCS/OH |
| Messor bouvieri | 1285 | -- | 0.58 (40) | -- | HCS/OH |
| Messor capitatus | 100 | -- | 1 (1) | -- | HCS/OH |
| Messor celiae | 11 | -- | -- | -- | HCS/OH |
| Messor lobicornis | 6 | -- | -- | -- | HCS/OH |
| Messor lusitanicus | 7 | -- | -- | -- | HCS/OH |
| Messor structor | 177 | -- | 1 (1) | -- | HCS/OH |
| Monomorium pharaonis | 50 | 0.33 (3) | -- | 17 | GO |
| Monomorium salomonis | 15 | -- | -- | -- | GO |
| Pheidole pallidula | 15655 | 0.13 (24) | 0 (2) | 1957 | GO |
| Plagiolepis pygmaea | 5416 | 0.21 (43) | 0 (19) | 1134 | GO |
| Plagiolepis schmitzii | 248 | 0.16 (31) | -- | 40 | GO |
| Proformica sp | 75 | -- | -- | -- | HCS/OH |
| Solenopsis sp | 47 | -- | -- | -- | C |
| Stenamma sp | 103 | -- | -- | -- | C |
| Tapinoma erraticum | 16033 | 1 (9) | 0 (1) | 16033 | GO |
| Tapinoma nigerimum | 26355 | 0.9 (148) | 0.93 (103) | 24040 | GO |
| Tapinoma simrothi | 136 | 1 (5) | -- | 136 | GO |
| Temnothorax sp | 138 | -- | -- | -- | C |
| Tetramorium sp | 3826 | -- | 0.17 (6) | -- | GO |
| Tetramorium sp2 | 1087 | -- | -- | -- | GO |
| Tetramorium sp3 | 127 | 0.2 (5) | -- | 25 | GO |

**Table S2**: Models testing the effect of the intensity of the ecological contrast on management differences. Linear mixed models showing variations between management regimes under different ecological contrast. Models include management, ecological contrast and their interaction as fixed factors, and locality and farm ID as random factors. In bold, significant models, with alpha = 0.1.

| **Response variable** | **ΔAIC _NULL_** | **X^2^** | **P** | **Marginal R^2^** | **Conditional R^2^** |
| --- | --- | --- | --- | --- | --- |
| **Ant abundance** | **1.59** | **7.58** | **0.055** | **5.1** | **29.0** |
| **Ant richness** | **6.10** | **12.11** | **0.007** | **8.7** | **26.3** |
| ***T.nigerrimum* abundance** | -4.4 | 1.60 | 0.660 | 1.8 | 33.4 |
| **APF (year)** | 0.45 | 5.55 | 0.136 | 3.6 | 24.2 |

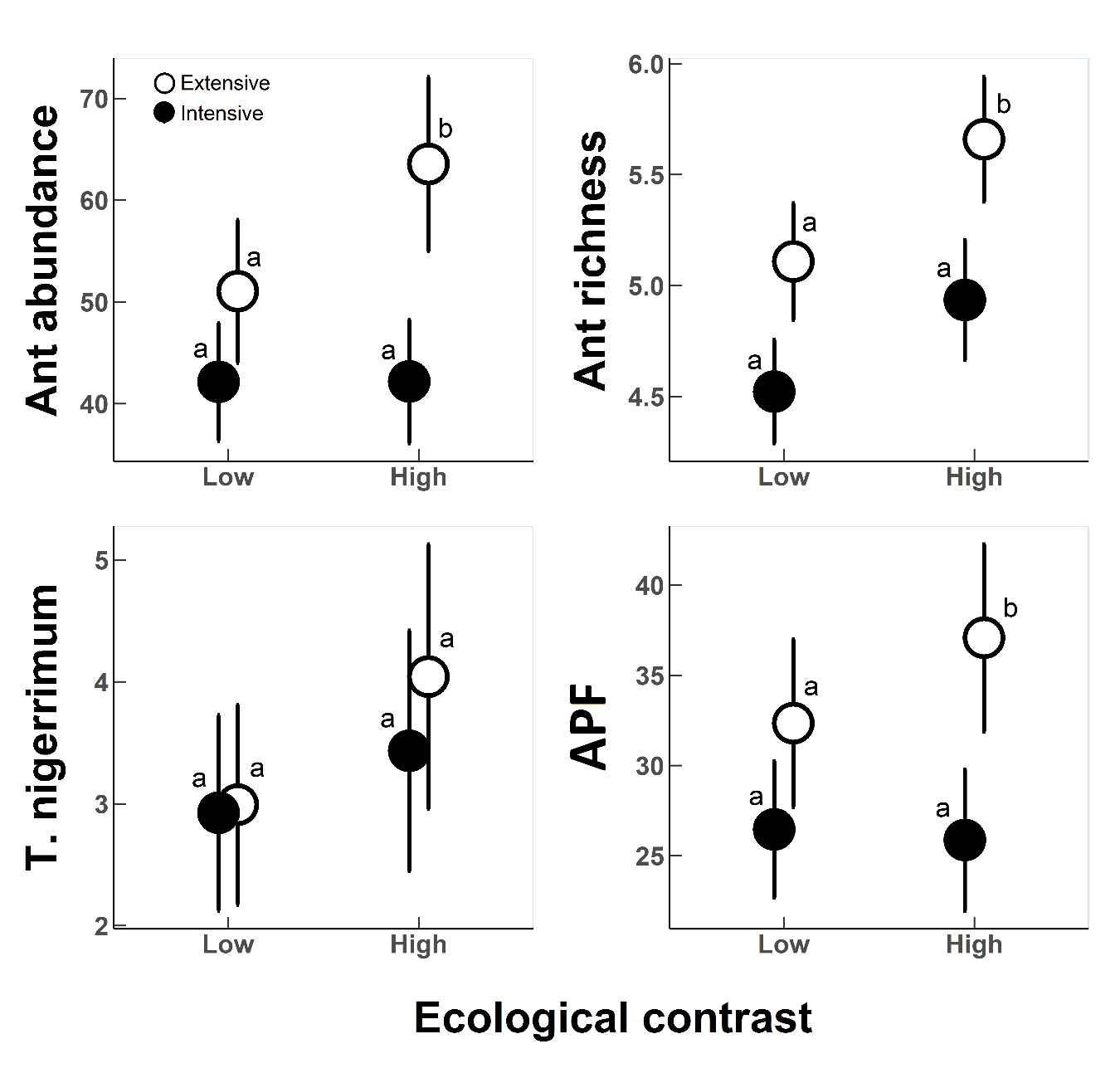

**Figure S1:** Effects of ecological contrast (Low contrast = Intensive Vs. extensive herb cover management; High contrast = Intensive Vs. Extensive plus organic -not pesticides- management) on ant abundance, ant richness, *T. nigerrimum* abundance and Ant Predation Function (APF) for the whole year. Letters show group assigned after Tukey post hoc comparison test (alpha=0.1) for differences in management at each level of ecological contrast).

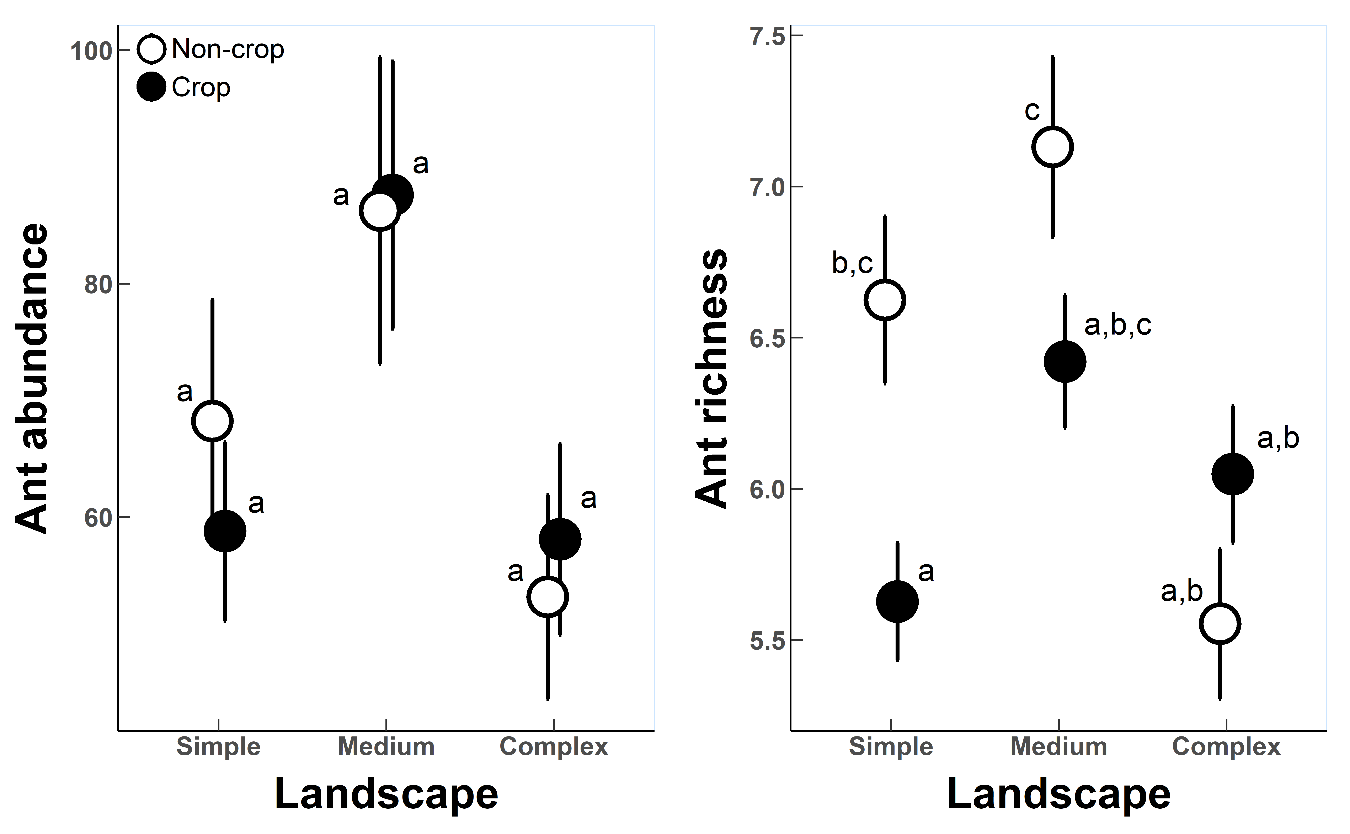

**Figure S2:** Differences in ant abundance and ant species richness across landscapes, for crop plots (black dots) and non-crop plots (white dots). Letters show Tukey post hoc multiple comparison test with an alpha=0.1 when statistically significant differences exist.

**References**

Roig & Spadaler (2010). Proposal of functional groups of ants for the Iberian Peninsula and Balearic Islands, and their use as bioindicators. *Iberomyrmex*, 2,2010.
